# Disentangling the evolutionary history of terrestrial planarians through phylogenomics

**DOI:** 10.1101/2025.01.07.631701

**Authors:** Lisandra Benítez-Álvarez, Nuria Escudero, Judit Salces-Ortiz, Iñaki Rojo, Fernando Ángel Fernández-Álvarez, Eduardo Mateos, Fernando Carbayo, Rosa Fernández

## Abstract

Triclads (Platyhelminthes, Tricladida) are found in marine, freshwater, and terrestrial habitats worldwide except Antarctica. Terrestrial planarians are grouped into the family Geoplanidae, which is subdivided into the subfamilies Geoplaninae, Bipaliinae, Rhynchodeminae, and Microplaninae. Some of these subfamilies result from taxonomic rearrangements based on molecular phylogenies inferred from a few molecular markers. However, the diagnosis of Rhynchodeminae was not aligned with the morphology of all its representatives. While the subfamilies are recovered as monophyletic in recent molecular phylogenies, robust hypotheses regarding the relationships between them remain unknown. In this study, we employ for the first time a phylogenomic framework to investigate the evolutionary relationships among the subfamilies, starting by obtaining the first transcriptomes for 15 species of terrestrial planarians. A total of 16 different datasets, comprising nearly two thousand single-copy genes inferred from transcriptomic data, were analyzed using various phylogenetic inference methods. We recovered, for the first time, a well-supported topology of phylogenetic relationships among Geoplanidae subfamilies, positioning Bipaliinae and Microplaninae as a clade sister to Rhynchodeminae + Geoplaninae. Internal relationships within the genus *Microplana* were not supported in our analyses. The subfamily Rhynchodeminae, represented in our phylogeny by species from the tribes Rhynchodemini and Caenoplanini, is re-diagnosed to align with previous taxonomic rearrangements. This study not only represents a significant step forward in the phylogenetic resolution of Geoplanidae but also provides important insights into the broader evolutionary dynamics shaping land planarian diversity.

**Highlights:**

- First transcriptomes of multiple terrestrial planarians sequenced.
- First molecular phylogeny of terrestrial planarians at subfamily level based on transcriptomic data.
- Highly supported topology for phylogenetic interrelationship among terrestrial planarians subfamilies.
- Unclear interrelationships between species from the genus *Microplana*.
- The subfamily Rhynchodeminae is re-diagnosed at the morphological level.

## 1. Introduction

Free-living flatworms belonging to the order Tricladida Lang, 1881 are represented in marine, freshwater and terrestrial habitats in all biogeographical areas of the world (Schockaert et al., 2008). The most diverse suborder, Continenticola Carranza et al., 1898, groups freshwater and terrestrial planarians in the superfamilies Geoplanoidea Stimpson, 1857 and Planarioidea Stimpson, 1857. Although the freshwater planarians are most known due to their high regeneration capabilities, as they are used as model organisms in research and academic fields (Newmark and Sánchez Alvarado, 2022; Saló and Baguñà, 2002; Sánchez Alvarado, 2006), land planarians (Geoplanidae Stimpson, 1857) have long intrigued researchers due to their diversity, evolutionary significance, and complex morphology, and in the last times due to the prevalence of invasive species with high impact in the economy and biodiversity of non-native areas (Alvarez-Presas et al., 2014; Lago-Barcia et al., 2019; Winsor et al., 2004). These organisms, found predominantly in tropical and subtropical regions, exhibit a variety of traits that make them suitable for studying patterns of speciation, biogeography, and adaptation (Álvarez-Presas et al., 2015, 2014; Alvarez-Presas et al., 2011).

The terrestrial planarians (Geoplanidae) are a monophyletic group classified into four subfamilies; Geoplaninae Stimpson, 1857, Bipaliinae Graff, 1896, Rhynchodeminae Graff, 1896, and Microplaninae Pantin, 1953 (Sluys et al., 2009) with a broad natural distribution across the world. The single exception is Geoplaninae, whose natural distribution is limited to the Central and South American regions. The internal phylogenetic relationships of the Geoplanidae have not yet been studied using a morphological approach, but for the Geoplaninae, the results produced non-robust trees that contradict molecular-based phylogenetic trees. This situation was seemingly caused by the limited number of valuable morphological characters and evolutionary events of convergence and reversal of character states (Lago-Barcia et al. 2023). Yet relationships within each of the subfamilies, based on Sanger-produced sequences are congruent with morphology in the sense that closely related species share a set of morphological characters (Alvarez-Presas et al., 2014, 2008; Álvarez-Presas and Riutort, 2014; Carbayo et al., 2013; Justine et al., 2024; Negrete et al., 2020; Riutort et al., 2012). These phylogenies were inferred from a limited number of molecular markers, such as nuclear, mitochondrial and ribosomal genes (see Álvarez-Presas and Riutort, 2014 for a comprehensive review). Different topologies have been reported in these studies, most of them showing low support values, where Bipaliinae has been recovered either as the earliest-splitting lineage in the clade of terrestrial planarians (Riutort et al., 2012) or as the sister group to Microplaninae (Álvarez-Presas and Riutort, 2014; Justine et al., 2024). Beyond the application of Sanger sequencing to understand land planarian interrelationships, mitogenomes were also sequenced for a few species representatives of the Geoplaninae (three species), Bipaliinae (six), and Rhynchodeminae (two) (Gastineau et al., 2020; Gastineau and Justine, 2020; Justine et al., 2024, 2020; Solà et al., 2015). Phylogenies using the full set of mitochondrial genes showed Geoplaninae as sister to Bipaliinae + Rhynchodeminae, albeit with low support. Therefore, the phylogenetic relationships between the subfamilies of terrestrial planarians remain unclear both based in Sanger-sequenced genes or mitogenomics (Gastineau and Justine, 2020; Justine et al., 2024, 2020).

Phylogenomics, which leverages genome-wide data to reconstruct evolutionary histories, offers a promising solution to these challenges. By providing greater amounts of phylogenetic signal accumulated in hundreds or thousands of genes, and reducing stochastic errors associated with individual gene trees, phylogenomic approaches can yield more robust and congruent topologies. Few studies have used transcriptome data to infer phylogenetic relationships within Platyhelminthes (Benítez-Álvarez et al., 2023; Egger et al., 2015; Laumer et al., 2015; Vila-Farré et al., 2023), but none of them included terrestrial planarians so far. In this study, we employ for the first time a phylogenomic framework to investigate the evolutionary relationships among land planarians, starting by obtaining the first transcriptomes of multiple terrestrial species (n=15). By utilizing high-throughput sequencing and comprehensive genomic data, we aim to overcome the limitations of earlier Sanger-based analyses and achieve a clearer understanding of their phylogeny. This study not only represents a significant step forward in the phylogenetic resolution of Geoplanidae but also provides important insights into the broader evolutionary dynamics shaping land planarian diversity.

## 2. Material and methods

### 2.1. Taxon sampling and sequencing

We hand-collected specimens of 15 terrestrial planarian species from Spain and Brazil. Detailed information on the sampled species, geographic coordinates, and localities is provided in Table S1. Samples were preserved in RNALater and stored at temperatures below -20°C until further processing. RNA extraction was conducted using the TRIzol® reagent (Invitrogen, USA) following the manufacturer’s protocol, with the addition of 5 µg of RNase-free glycogen as a carrier for low-input samples in the aqueous phase. RNA concentrations were measured using the Qubit RNA BR Assay kit (Thermo Fisher Scientific). Libraries were prepared with Illumina’s TruSeq Stranded mRNA kit and sequenced on a NovaSeq 6000 platform (Illumina, 2 × 150 bp) to achieve 6Gb coverage.

### 2.2. Phylogenetic inference

We obtained a dataset of 18 species representative of the Geoplanoidea superfamily, including 3 freshwater species belonging to the Dugesiidae family and the newly-generated 15 terrestrial species belonging to Geoplanidae (Table S1). All of them resulted in a high BUSCO completeness (>80%) of metazoa database, which seems to be common for Platyhelminthes (Martínez-Redondo et al., 2024).

*De novo* transcriptome assemblies and longest isoforms were obtained following the pipeline described in MateDB2 database (Martínez-Redondo et al., 2024). The longest isoforms were analyzed with OrthoFinder2 (Emms and Kelly, 2019, 2015) to infer orthologous groups. We retrieved all single copy orthologous groups (SC hereafter; n=1873) both at the amino acid (Dataset 1) and nucleotide (Dataset 2) level for further analysis. Additionally, from these 1873 single copy orthogroup dataset, we removed 1% and 10% of outliers with genesortR (Koch, 2021) (Datasets 3-6). Afterwards, we retained the 75% and the 50% of top genes, to obtain the datasets 7 to 14. Finally, we built the datasets 15 and 16 including only *Microplana* species, in order to further explore the phylogenetic relationships within the genus (Table 1).

**Table 1.** Datasets used in phylogenetic analyses. SC: Single Copy genes, MSC: Multispecies Coalescent model (ASTRAL-Pro), ML: Maximum Likelihood (IQ-TREE2). All datasets contain 100% of species completeness.

| Dataset | Number of species | Number of SC | Data type | Phylogenetic inference method | Description |
| --- | --- | --- | --- | --- | --- |
| Dataset 1 | 18 | 1873 | protein | MSC / ML | SC genes from Orthofinder |
| Dataset 2 | 18 | 1873 | nucleotide | MSC / ML | SC Copy genes from Orthofinder |
| Dataset 3 | 18 | 1855 | protein | MSC | Deleting 1% of outliers from dataset 1 |
| Dataset 4 | 18 | 1855 | nucleotide | MSC | Deleting 1% of outliers from dataset 2 |
| Dataset 5 | 18 | 1686 | protein | MSC/ML | Deleting 10% of outliers from dataset 1 |
| Dataset 6 | 18 | 1686 | nucleotide | MSC/ML | Deleting 10% of outliers from dataset 2 |
| Dataset 7 | 18 | 1391 | protein | MSC | 75% top of deleting 1% of outliers from dataset 1 |
| Dataset 8 | 18 | 1391 | nucleotide | MSC | 75% top of deleting 1% of outliers from dataset 2 |
| Dataset 9 | 18 | 1265 | protein | MSC | 75% top of deleting 10% of outliers from dataset 1 |
| Dataset 10 | 18 | 1265 | nucleotide | MSC | 75% top of deleting 10% of outliers from dataset 2 |
| Dataset 11 | 18 | 927 | protein | MSC/ML | 50% top of deleting 1% of outliers from dataset 1 |
| Dataset 12 | 18 | 927 | nucleotide | MSC/ML | 50% top of deleting 1% of outliers from dataset 2 |
| Dataset 13 | 18 | 843 | protein | MSC/ML | 50% top of deleting 10% of outliers from dataset 1 |
| Dataset 14 | 18 | 843 | nucleotide | MSC/ML | 50% top of deleting 10% of outliers from dataset 2 |
| Dataset 15 | 5 | 843 | protein | MSC/ML | 50% top of deleting 10% of outliers from dataset 1 only *Microplana* sp. |
| Dataset 16 | 5 | 843 | nucleotide | MSC/ML | 50% top of deleting 10% of outliers from dataset 2 only *Microplana* sp. |

Orthogroups in each dataset were independently processed using PREQUAL (Whelan et al., 2018) to mask regions with non-homologous characters, and subsequently aligned with MAFFT with auto option and 1000 iterative refinement. Finally, the alignments were trimmed to remove poorly aligned regions with trimAL using the ‘–automated1’ flag (Capella-Gutiérrez et al., 2009). The gene trees were obtained using Maximum Likelihood (ML), as implemented in IQ-TREE2 (Minh et al., 2020), with the mixture model strategy described below for nucleotide and protein data, and 10,000 ultrafast bootstrap replicates.

Species trees for all datasets were obtained from gene trees using the Multispecies Coalescent method (MSC) implemented in ASTRAL-Pro 2 (Zhang and Mirarab, 2022) with default options. In the case of analyses performed with datasets 15 and 16, the -t2 option implemented in ASTRAL-Pro 2 (Zhang and Mirarab, 2022) was used to recover the full branch annotation. The obtained information was summarized in pie charts using the AstralPlane package (Hutter, 2024) in R.

A supermatrix approach was explored for datasets 1-2, 5-6, 11-14 which showed higher final normalized quartet scores (FNQS) and branch support with MSC method. (Table 1, Table S2). The ML trees for datasets were obtained from concatenated matrices using mixture models. For protein data, we used the trees obtained with MSC as starting tree and the LG model with 20 categories (C20), Gamma rate heterogeneity calculation (+G), site-specific frequency profile inference (+F), and 10,000 ultrafast bootstrap replicates and 10,000 replicates for SH-like approximate likelihood ratio (SH-aLRT) test. On the other hand, nucleotide sequences were analyzed using the MIX option with three components (JC, HKY, and GTR), four Gamma categories (+G4), and 10,000 replicates for ultrafast bootstrap and SH-aLRT test.

## 3. Results

### A phylogenomic hypothesis for the interrelationships of land planarians

Several phylogenetic analyses were conducted to infer the phylogeny of the terrestrial planarians included in our study (Table 1). All nucleotide data matrices recovered two strongly supported clades: one including the Rhynchodeminae and Geoplaninae subfamilies, and another including the Bipaliinae and Microplaninae subfamilies (Fig. 1). However, some matrices analyzed at the amino acid level yielded either low support or supported alternative topologies (Appendix A and B).

**Fig. 1.**
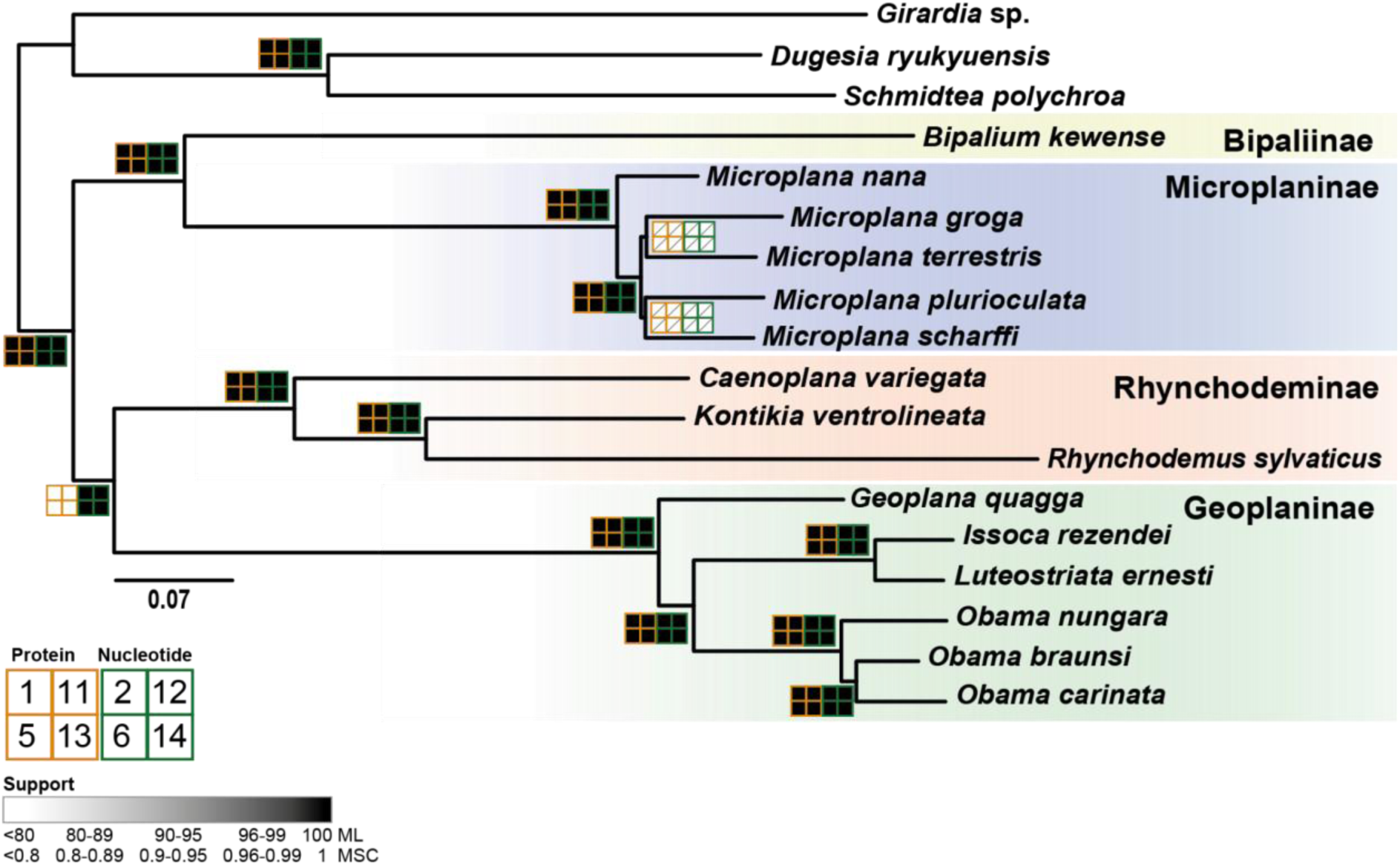
Summary of phylogenetic relationships obtained with the different datasets. The support values obtained with both inference methods (Maximum Likelihood (ML) and Multispecies Coalescent Model (MSM)) are indicated in a grid on the nodes for datasets 1-2, 5-6 and 11-14. Each grid represents a dataset. White square indicates low support (< 80/0.8) and slash inside indicates an alternative hypothesis recovered for that node (see Fig. 2 and Appendices A-C). The support values are indicated by a scale colour from white (<80/0.8) to black (100/1). Scale bar: substitutions per site. See Table 1 for more information about datasets and Appendix A and B for the inferred topologies of all analyses.

Sister relationships with high support values, based on nucleotide data, included Rhynchodeminae + Geoplaninae and Bipaliinae + Microplaninae. Since the protein data was insufficient to resolve the internal relationships within the family Geoplanidae, we focus our discussion on the results obtained from the nucleotide data. High support values were consistently observed across all analyses for the internal relationships within the subfamily Geoplaninae (Fig. 1). *Geoplana quagga* Marcus, 1951 represent the first lineage to diverge as the sister group of the clade formed by *Obama* Carbayo, Álvarez-Presas, Olivares, Marques, E.M. Froehlich & Riutort, 2013 and the sister genera *Issoca* C.G. Froehlich, 1954 and *Luteostriata* Carbayo, 2010. In the same way, *Caenoplana* Moseley, 1877 was recovered as sister to *Kontikia* C.G. Froehlich, 1954 and *Rhynchodemus* Leidy, 1852 in the subfamily Rhynchodeminae.

On the other hand, the internal relationships of the subfamily Microplaninae resulted in different interrelationships depending on the analyses. The only position that was supported with all datasets (1-14) was the sister relationship of *Microplana nana* Mateos, Giribet & Carranza, 1998 to the clade formed by *Microplana groga* Jones, Webster, Littlewood & McDonald, 2008, *Microplana plurioculata* Sluys, Mateos & Álvarez-Presas, 2016*, Microplana scharffi* (Von Graff, 1896), and *Microplana terrestris* (Müller 1774).

To further explore the phylogenetic relationships within this genus, we inferred phylogenetic relationships by constraining *M. nana* as sister to the other *Microplana* species, as supported by the analyses described above (Fig. 2).

**Fig. 2.**
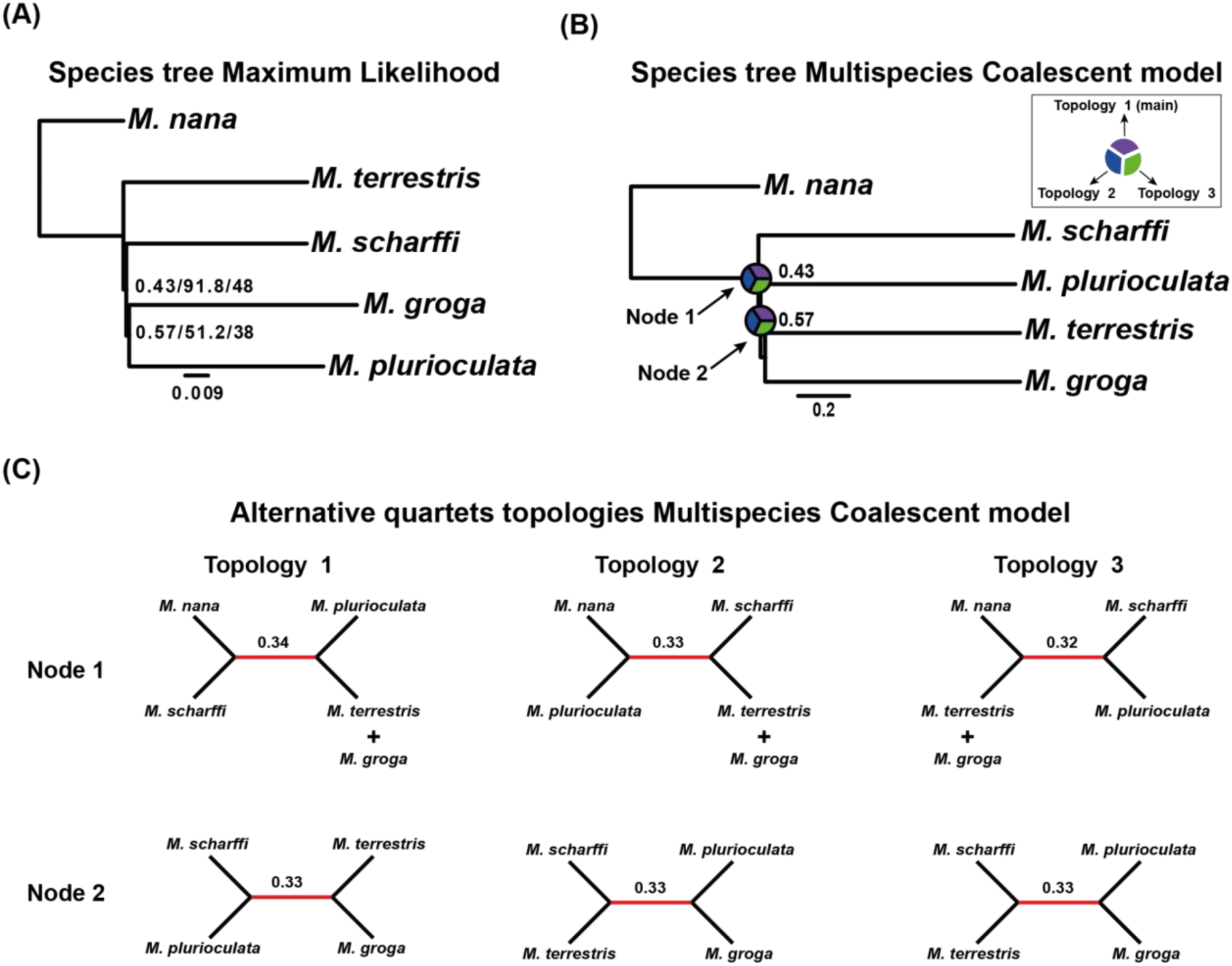
Phylogeny between *Microplana* species. Species trees obtained analyzing dataset 16 including nucleotide sequences of 843 single copy genes and only *Microplana* representatives. (A): Maximum Likelihood tree. Values in the nodes correspond to branch support estimated for mixture model/support for SH-aLRT test/ultrafast bootstrap. Scale bar: substitutions per site; (B): Multispecies Coalescent model implemented in ASTRAL-Pro. Pie charts in the nodes indicate the per branch quartet support for the main (topology 1) and the alternative topologies (topology 2 and 3). Scale bar. coalescent units. (C): Alternative topologies with the corresponding quartet support for the two nodes estimated in ASTRAL-Pro.

The short branch lengths in the tree shown in Fig. 2 and the similar quartet support of the alternative topology indicate high discordance among gene trees with the consequent low resolution in the species tree topology for the Microplaninae. In the same way, low bootstrap support values were found in the Maximum Likelihood trees (Appendix C).

The very close per branch quartet support values of all possible quartet topologies around nodes 1 and 2 in the tree (Fig. 2 B) could indicate an incomplete lineage sorting process in these species. This may be supported by the similar per branch quartet support values for the main and the alternative topologies. Additionally, the short genetic distances among them indicated by both, the coalescent units in the case of trees obtained using MSC and the substitutions per site in the case of ML trees (Appendix C) indicate a high incongruence in the multilocus data.

## 4. Discussion

Here, we obtained the first phylogeny based on transcriptomic data of terrestrial planarians making available 15 transcriptomes of representative species of this lineage, including the invasive species *Obama nungara* and *Bipalium kewense*. A strongly supported topology among the four subfamilies of land planarians was recovered in our study. We found that Bipaliinae and Microplaninae form a sister-group relationship, and together they are sister to a clade comprising Rhynchodeminae and Geoplaninae. By contrast, earlier works placed Rhynchodeminae as the earliest-splitting lineage among terrestrial planarians (Álvarez-Presas and Riutort, 2014), while other analyses recovered Bipaliinae as sister to a clade uniting Microplaninae, Rhynchodeminae, and Geoplaninae (Alvarez-Presas et al., 2008; Solà et al., 2023). Additionally, mitogenome-based studies suggested yet another arrangement, with Geoplaninae as the external group to a clade containing Rhynchodeminae, Bipaliinae, and Microplaninae (Gastineu et al., 2024). Within Geoplaninae—the subfamily most extensively represented in our dataset—internal relationships were highly supported, aligning with previous findings in a taxonomic revision of the Geoplaninae clade (Fernando Carbayo et al., 2013). Nonetheless, it is noteworthy that many of these analyses, including ours, show low support at certain basal nodes, underscoring the need for caution in interpreting these topologies.

Our phylogenies represent a step forward compared to previous works, as a robust hypothesis for the evolutionary relationships between the higher Geoplanidae taxa has finally emerged. According to this hypothesis, the Bipaliinae are sister to the Microplaninae, while the Rhynchodeminae are sister to the Geoplaninae, with all four taxa forming a monophyletic group. Biogeographically, there is no evidence of close relationships, as the natural distribution of one subfamily of each clade is restricted to a specific region (e.g., Geoplaninae in the neotropics; Bipaliinae in Madagascar, and Southern, Eastern, and Southeast Asia), while the other member of each pair is cosmopolitan. Remarkably, there is no morphological evidence supporting the topology of our tree, nor any alternative one. This could be due to (a) the diagnostic features of the subfamilies being currently difficult to compare (see Table S3) and (b) the diagnoses of the subfamilies not aligning with later taxonomic rearrangements.

Comparisons between taxa require detailed species descriptions. However, old descriptions do not always conform to current standards, and therefore, relevant morphological details from many of them remain unknown, particularly those of the subepidermal and parenchymal musculature and the copulatory apparatus (Ogren and Kawakatsu, 1991). This situation hinders the search for morphological synapomorphies supporting each pair of subfamilies based solely on the literature. Alternatively, the absence of synapomorphies might be due to the ancient origin of the Geoplanidae. While the origin of the Geoplanidae remains uncertain, that of the Bipaliinae has been molecularly estimated to be between 395.4 and 123.63 million years ago (Solà et al., 2023). This ancient origin likely allowed convergence and character state reversal to occur, as suggested for the Geoplaninae (Lago-Barcia et al., 2023), thus blurring morphological evidence of relationships within the group.

Progress in understanding the phylogeny of the Tricladida and Geoplanidae has been achieved through molecular-based approaches, notably (Baguñà et al., 2001) and (Alvarez-Presas et al., 2008). Based on this new perspective, some taxonomic changes were introduced into the classification system of land flatworms (Sluys et al., 2009). Of particular interest here is the repositioning of the multi-eyed rhynchodemin tribes Pelmatoplanini Ogren & Kawakatsu, 1991 and Caenoplanini Ogren & Kawakatsu, 1991 (Sluys et al., 2009). However, the diagnosis of Rhynchodeminae was not updated accordingly. Therefore, we propose to amend the diagnosis of the Rhynchodeminae to align with the morphology of its ingroups, particularly with respect to the eyes, the width of the creeping sole, and cephalic specializations (Table S3), as follows:

*Re-diagnosis of the Subfamily Rhynchodeminae Graff, 1896.* Geoplanidae of elongate cylindroid form with two eyes near the simple, tapered anterior end or with multiple eyes around the anterior end, generally continuing posteriorly; no tentacles or headplate; with well-defined creeping sole, occupying 25-84% of the ventral surface. Anterior extremity may have a sucker on the ventral surface (*Cotyloplana*) or a cephalic retractor muscle (*Pimea*). Ventral testes, sometimes dorso-ventral (Anzoplanini; *Marionfyfea*).

Regarding the subfamily Microplaninae, this lineage is considered endemic in Europe, although some species have been found on the African (Jones, 1998), American (Ogren and Kawakatsu, 1998; Murchie and Justine, 2021) and Asian continents (Kawakatsu and Ogren, 1998). All studies conducted on them highlighted their high species richness (Alvarez-Presas et al., 2022; Jones et al., 2008; Jones and McDonald, 2021; Vila-Farré et al., 2011), estimating that many of them still remain to be discovered (Mateos et al., 2017). The phylogenetic position of Microplaninae has been re-evaluated several times using molecular data from mitochondrial, ribosomal and nuclear genes (cox1, 18S, 28S and EF1α), with different results depending on the study. Microplaninae has been closely associated with Geoplaninae (Jones, 1998), Rhynchodeminae (Alvarez-Presas et al., 2008) and Bipaliinae (Álvarez-Presas and Riutort, 2014) by different authors. In this study, we show a highly supported relationship between Microplaninae and Bipaliinae. However, the internal relationships inside this subfamily remain obscure.

Despite the analyses of hundreds of loci from our newly generated transcriptomic datasets, the interrelationships between *Microplana* species remain unresolved. The only supported relationship is the position of *M. nana* as the sister group of the remaining studied *Microplana* species as shown in previous molecular analyses (Mateos et al., 2017; Sluys et al., 2016). However, in these analyses no support was obtained for the relationships among other *Microplana* species and low genetic distances among them were reported (Mateos et al., 2017; Sluys et al., 2016). Therefore, although morphological data could add valuable information for species identification and delimitation in integrative taxonomic approaches, the evolutionary history of this lineage remains unclear (Sluys et al., 2016). Phylogenetic inference methods can be masked by biological processes as hybridization or incomplete lineage sorting (ILS) (Pezzi et al., 2024; Wang et al., 2018). Although MSC implemented in ASTRAL-Pro theoretically can handle ILS with high efficiency (Liu et al., 2015; Mirarab et al., 2016), complex evolutionary histories can escape the accuracy of this method. Therefore, the population dynamics linked to biological and adaptive processes can affect the species tree estimation. Biogeographical events and adaptive response to similar habitat conditions could underlay a complex evolutionary history challenging to infer with the data and methods currently at hand. Hence, more extensive studies—including additional species and datasets—will be needed to clarify *Microplana*’s complex evolutionary history.

## 5. Final remarks

Our transcriptome-based approach has significantly advanced the understanding of terrestrial planarian phylogeny, providing robust support for the relationships among the four geoplanid subfamilies while also prompting a needed re-diagnosis of Rhynchodeminae. Nonetheless, as evidenced by the unresolved internal relationships within *Microplana*, challenges remain in fully disentangling the evolutionary history of these ancient lineages. Incomplete lineage sorting, potential hybridization, and limited morphological data—especially from older descriptions—complicate phylogenetic reconstruction. Moreover, although previous studies have suggested the inclusion of certain freshwater species (*Reynoldsonia* Ball, 1974; *Spathula* Nurse, 1950; and *Romankenkius* Ball, 1974) within the terrestrial clade, implying a possible return to freshwater habitats after terrestrialization (Alvarez-Presas et al., 2008; Álvarez-Presas and Riutort, 2014), our study focused solely on terrestrial representatives. Future research incorporating these understudied lineages, alongside broader species sampling and refined morphological analyses, will be key for clarifying the deep divergences and potential habitat shifts within the group.

Altogether, this work underscores both the promise and the complexity of unraveling the evolutionary chronicle of land planarians. By illuminating key phylogenetic relationships and highlighting gaps in our current knowledge, we pave the way for integrative taxonomic efforts and further genomic studies that will ultimately deepen our understanding of the evolutionary processes shaping these remarkable organisms.

## Declaration of Generative AI and AI-assisted technologies in the writing process

All text was human written, with some parts AI-refined.

## CRediT authorship contribution statement

Conceptualization: LB-A, FC, RF; Methodology: LB-A, NE, JS-O, RF; Software: LB-A; Formal analysis: LB-A; Investigation: LB-A, NE, JS-O; Resources: IR, FAFA; Writing - original draft: LB-A, FC, RF; Writing - review and editing: all authors; Visualization: LB-A; Supervision: RF; Funding acquisition: FC, RF

## Acknowledgements

RF acknowledges support from the following sources of funding: Ramón y Cajal fellowship (grant agreement no. RYC2017-22492 funded by MCIN/AEI 10.13039/501100011033 and ESF ‘Investing in your future’), the European Research Council (this project has received funding from the European Research Council (ERC) under the European’s Union’s Horizon 2020 research and innovation programme (grant agreement no. 948281)) and the Secretaria d’Universitats i Recerca del Departament d’Economia i Coneixement de la Generalitat de Catalunya (AGAUR 2021-SGR00420). We also thank Centro de Supercomputación de Galicia and CSIC for access to computer resources (CESGA and DRAGO respectively).

FÁF-Á was supported by a Beatriu de Pinós fellowship from Secretaria d’Universitats i Recerca del Departament de Recerca i Universitats of the Generalitat de Catalunya (Ref. BP 2021 00035) and a Ramón y Cajal fellowship (Ref. RYC2023-043494-I) funded by MCIN/AEI 10.13039/501100011033 and FSE+. This research was supported by the Spanish government through the ‘Severo Ochoa Centre of Excellence’ accreditation (CEX2019-000928-S).

## Data accessibility statement

Intermediate files and outputs for reproducibility of analyses are accessible at: https://github.com/MetazoaPhylogenomicsLab/Benitez-Alvarez_et_al_2025_Phylogenomics_land_planarians

---

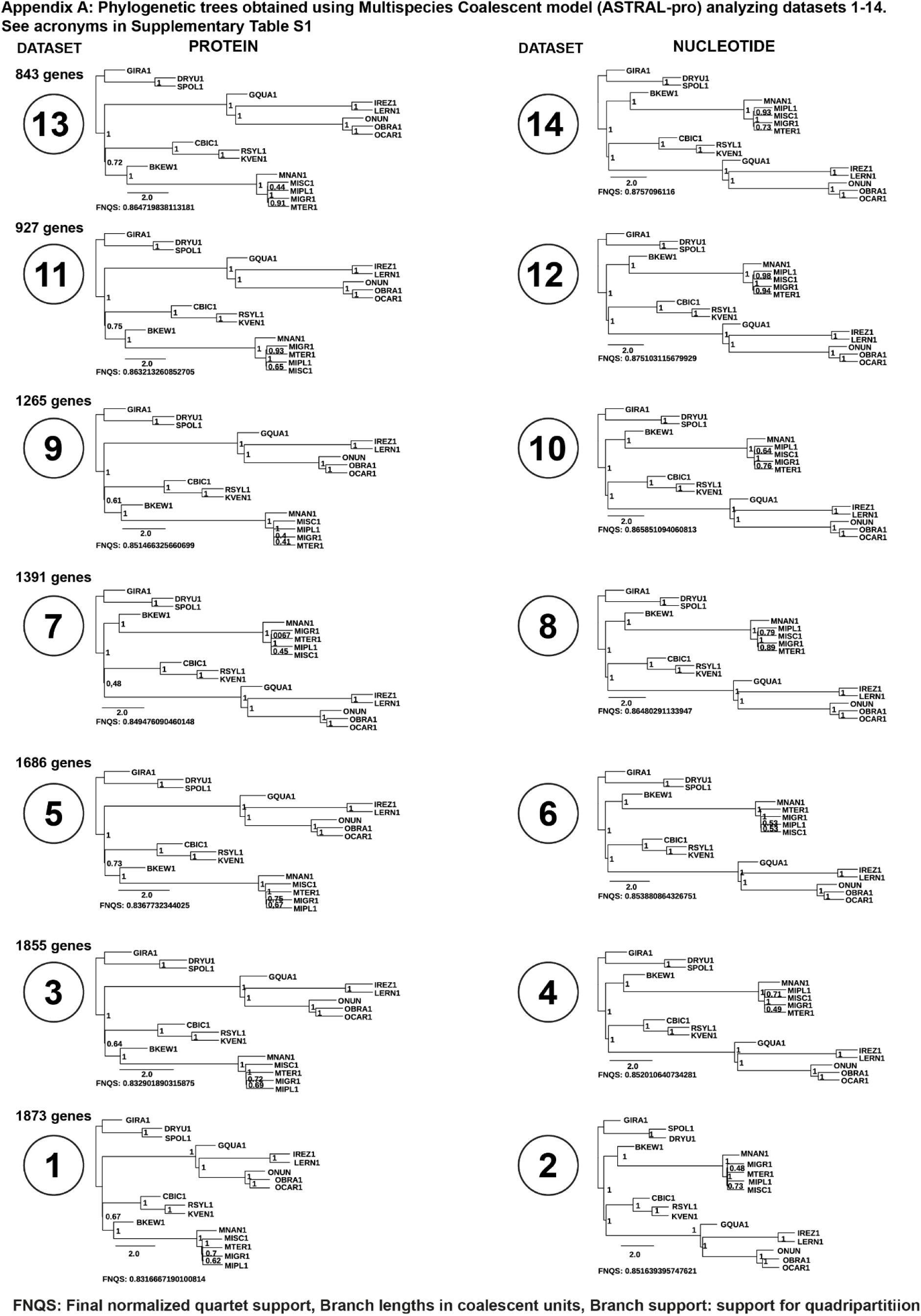

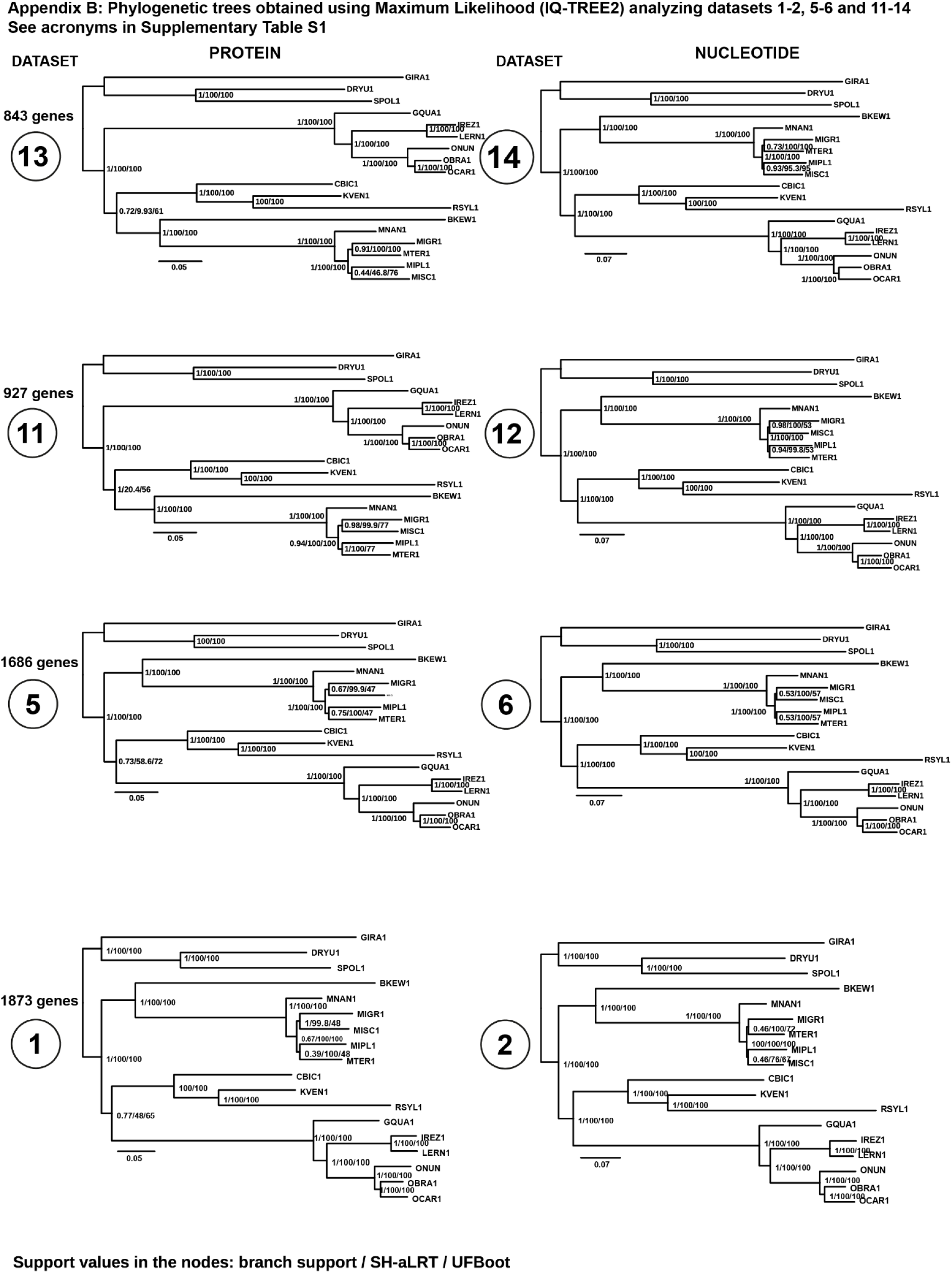

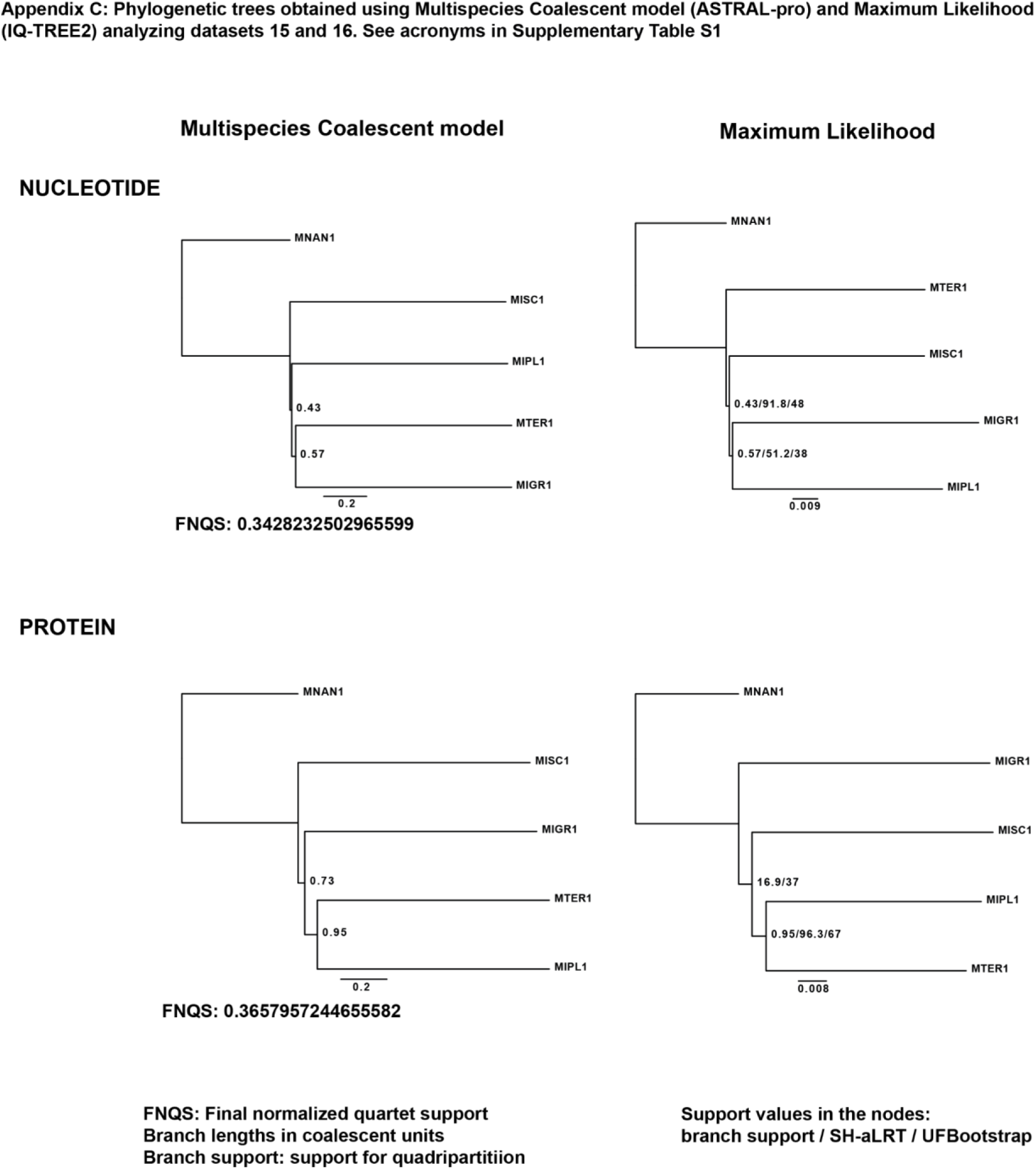

